# A DNA repair-independent role for alkyladenine DNA glycosylase in alkylation-induced unfolded protein response

**DOI:** 10.1101/2021.03.31.437844

**Authors:** Larissa Milano, Clara F. Charlier, Rafaela Andreguetti, Thomas Cox, Eleanor Healing, Marcos P. Thomé, Ruan M. Elliott, Leona D. Samson, Jean-Yves Masson, Guido Lenz, João Antonio P. Henriques, Axel Nohturfft, Lisiane B. Meira

## Abstract

Alkylating agents damage DNA and proteins and are widely used in cancer chemotherapy. While the cellular responses to alkylation-induced DNA damage have been explored, knowledge of how alkylation damage affects global cellular stress responses is still sparse. Here, we examined the effects of the alkylating agent methylmethane sulfonate (MMS) on gene expression in mouse liver taking advantage of mice deficient in alkyladenine DNA glycosylase (Aag), the enzyme that initiates the repair of alkylated DNA bases. MMS induced a robust transcriptional response in wild-type liver that included markers of the endoplasmic reticulum (ER) stress/unfolded protein response (UPR) known to be controlled by the transcription factor XBP1, a key UPR effector. Importantly, this response is significantly reduced in the *Aag* knockout. To investigate a potential role for AAG in alkylation-induced UPR, the expression of UPR markers after MMS treatment was interrogated in human glioblastoma cell lines expressing different AAG levels. Alkylation induced the UPR in cells expressing AAG; conversely, *AAG* knock-down compromised UPR induction and led to a defect in XBP1 activation plus a decrease in the expression of the ER chaperone BiP. To verify that the DNA repair activity of AAG is required for this response, *AAG* knockdown cells were complemented with wild-type *Aag* or with a mutant version of the *Aag* gene producing a glycosylase-deficient AAG protein. As expected, the glycosylase-defective mutant Aag does not fully protect *AAG* knockdown cells against MMS-induced cytotoxicity. Remarkably, however, alkylation-induced XBP1 activation is fully complemented by the catalytically inactive AAG enzyme. This work establishes that, in addition to its enzymatic activity, AAG has non-canonical functions in alkylation-induced UPR that contribute to the overall cellular response to alkylation.

**Significance Statement:** Stress response pathways, such as the DNA damage response (DDR) and the UPR, are critical in both the etiology and treatment of cancer and other chronic diseases. Knowledge of an interplay between ER stress and genome damage repair is emerging, but evidence linking defective DNA repair and impaired ER stress response is lacking. Here, we show that AAG is necessary for UPR activation in response to alkylating agents. AAG-deficient mice and human cancer cells are impaired in alkylation-induced UPR. Strikingly, this defect can be complemented by an AAG variant defective in glycosylase activity. Our studies suggest AAG has non-canonical functions and identify AAG as a point of convergence for stress response pathways. This knowledge could be explored to improve cancer treatment.

## Introduction

Organisms are constantly exposed to a variety of stresses that can result in macromolecular injury and cellular dysfunction (1, 2). Reactive compounds that originate from the environment or arise from intracellular processes can damage nucleic acids, proteins and lipids. Exposure to stress triggers highly conserved signalling pathways, such as the DNA damage response (DDR) and ER stress responses that act to restore homeostasis. Failure of cells and tissues to properly respond to stress and damage underpins the pathogenesis of many diseases (2, 3).

Alkylating agents represent an abundant and ubiquitous family of reactive chemicals that can damage DNA, RNA and proteins (4). Sources of alkylating agents include by-products of metabolism (5), and environmental nitroso-compounds such as nitrosamines that are present in pollutants, food preservatives and recently also found in common medications (6–8). Exposure to alkylating agents has been associated with cancer, type-2 diabetes, non-alcoholic steatohepatitis and neurodegenerative diseases (9, 10). On the other hand, because they effectively kill dividing tumour cells, alkylating agents are commonly employed as cancer chemotherapeutic agents.

Alkylation-induced DNA base lesions are primarily repaired by the base excision repair (BER) pathway, initiated by the enzyme alkyladenine DNA glycosylase, or AAG (also known as MPG) (5). AAG excises damaged bases from the DNA phosphodiester backbone, generating an abasic site. Subsequently, an apurinic/apyrimidinic endonuclease cleaves the phosphodiester backbone at the abasic site, creating a single-strand break (SSB) that contains 3’OH and 5’deoxyribose-5-phosphate (5’dRP) termini. Next, DNA polymerase β removes the 5’dRP and carries out single-nucleotide gap filling synthesis. The nicked DNA is then ligated by DNA ligase I or the XRCC1/Ligase III complex (5, 11). The flux of intermediates through this pathway must be efficiently coordinated because BER intermediates, such as abasic sites and SSB, are toxic (12–14). It has been shown that BER flux imbalance due to AAG overexpression exacerbates alkylation toxicity (15, 16). Moreover, elevated AAG expression has been associated with poor prognosis in patients with glioblastoma, an aggressive type of brain cancer often treated with alkylating agents (17, 18).

While the effects of alkylation on DNA have been well studied, cellular responses to alkylation-induced protein damage are still poorly understood. Alkylation treatment of yeast and mammalian cells induces hallmarks of ER stress, involving the UPR (19–21). The UPR is an adaptive signal transduction pathway orchestrated by the ER that is important for the maintenance of a functional proteome. A wide range of perturbations can result in ER stress, such as accumulation of unfolded/misfolded proteins, disturbances in calcium homeostasis, hypoxia, oxidative stress and viral infections (22). Three ER-resident transmembrane proteins initiate the UPR: inositol-requiring kinase 1 (IRE1), activating transcription factor 6 (ATF6) and protein kinase-like ER kinase (PERK). These transducers are negatively regulated by chaperones that dissociate during ER stress, leading to activation of the UPR (22). The UPR usually acts as a cytoprotective mechanism, but chronic ER stress leads to cell death (22). UPR activation was shown to play a key role in cancer, by enabling tumour cells to tolerate and thrive in a hostile environment of nutrient deprivation, hypoxia and low pH, which in turn contributes to cellular transformation, tumour growth, metastasis and resistance to chemo/radiotherapy (23).

AAG was shown to be the major DNA glycosylase activity for the excision of alkylated lesions in mouse liver (24). To better characterize the outcomes of alkylation damage, we analyzed gene expression in livers of wild-type and Aag-deficient mice that had been exposed to the model alkylating agent MMS. Our findings show that alkylation treatment induces ER stress and the UPR *in vivo*. Surprisingly, alkylation-induced expression of ER stress genes was significantly reduced in *Aag*^−/−^ livers. To probe the mechanism underlying this relationship, we employed a panel of human glioblastoma cell lines expressing different levels of AAG and DNA repair proficient and deficient mouse AAG variants. As in mouse liver, we find AAG is required for optimal UPR induction after alkylation treatment in human glioblastoma cells. Moreover, we show that a repair-defective AAG variant unable to complement alkylation-induced PARP activation and survival is proficient in the activation of the bZIP transcription factor X-box binding protein 1 (XBP1), a key regulator of the UPR. This striking result indicates AAG has non-canonical roles in alkylation-induced UPR induction. Finally, considering the dual survival or cell death outcome of the UPR, we also examine the effect of AAG status on cellular sensitivity to alkylating agents alone or combined with a pharmacological activator of ER stress.

## Results

### Aag is required for alkylation-induced ER stress response mediated by Xbp1 activation

Alkylating agents activate both the DDR and the UPR. To explore potential connections between these pathways, we compared alkylation-induced changes in livers of wild-type and Aag-deficient mice. Wild-type and *Aag*^−/−^ animals were injected with a mild, non-lethal dose of the direct acting alkylating agent MMS, and liver tissue was harvested after 6 hours. This MMS dose and time point have been previously characterised, and for both genotypes no difference in liver histology or weights between controls or MMS-treated cohorts was noted by pathological examination (16).

Following RNA extraction, transcriptome analysis was performed using gene chip arrays. Changes in expression associated with a fold change of at least 1.75 (false discovery rate-adjusted p value ≤ 0.05) were considered as significant, and the affected genes marked for further analysis. Only minor differences in gene expression were observed under control conditions between wild-type and Aag-deficient liver, indicating that the absence of Aag does not cause significant stress under basal metabolic conditions (Fig. 1A, Fig. S1).

**Fig. 1.**
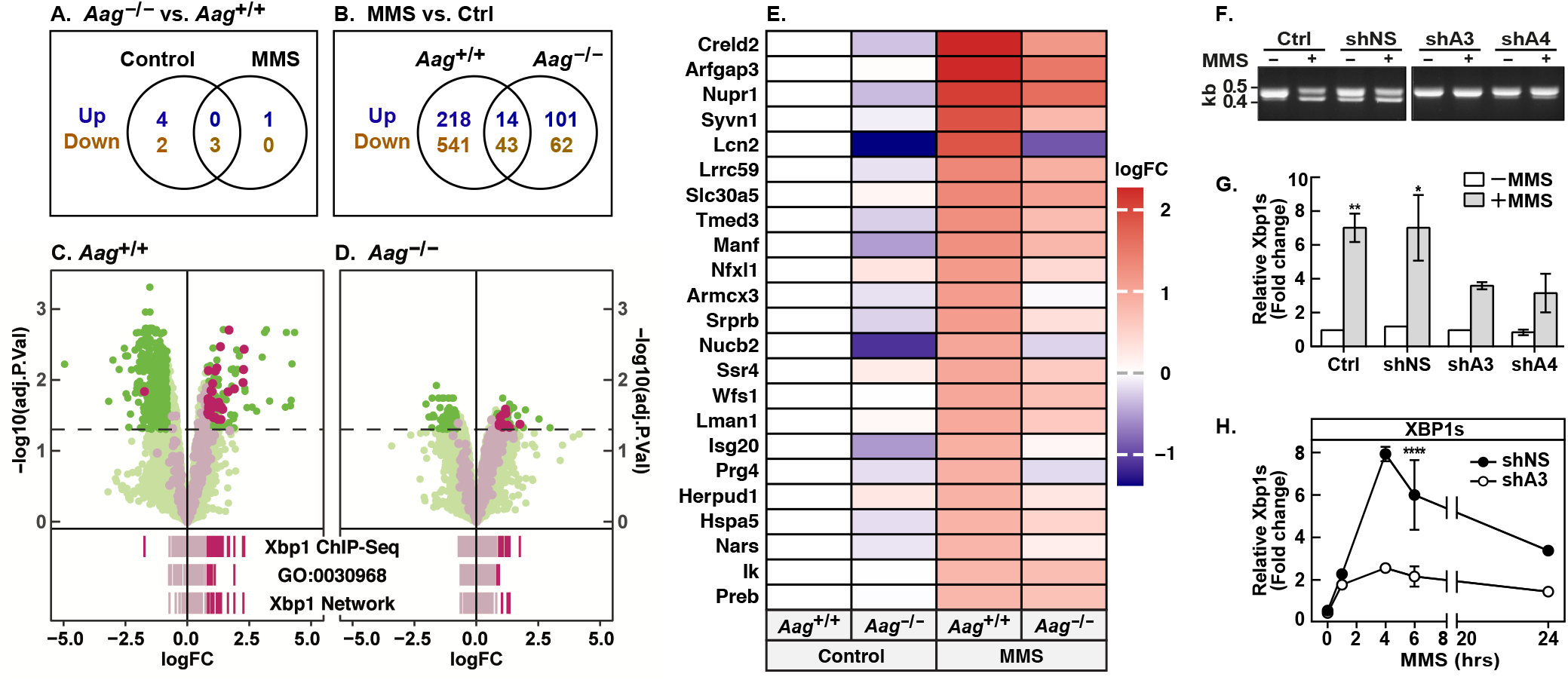
Aag modifies the transcriptional response to alkylation and is required for XBP1 splicing induced by alkylation. Wild-type and Aag-deficient mice (n=3) were injected with MMS or solvent and euthanised 6 h later. Liver RNA was analyzed using oligonucleotide microarrays. (A and B) Venn diagrams indicating number of differentially regulated probe sets (log_2_ fold change ≥ 1.75; false discovery rate (FDR)-adjusted p value ≤ 0.05). Detailed gene expression data are given in Supplemental Data 1. (C and D) Negative log_10_ adjusted p values are plotted against log2 (fold change). Dashed line, negative log_10_ (0.05). Xbp1 targets according to mouse liver ChIP-seq data (29) are highlighted in gray or in magenta where |log_2_ (fold change)| ≥ 1.75 and p-FDR ≤ 0.05. Rug plots below indicate the log_2_ (fold change) of genes annotated as Xbp1 targets (“Xbp1 ChpSq”), ER stress response (“GO:0030968”) or Xbp1 transcriptional correlation network (“Xbp1 Netwrk”); where |log_2_ (fold change)| < 1.75 genes are marked in gray; lines are drawn at 60% transparency. (E) Heatmap indicating log_2_ fold changes of known Xbp1 target genes (28) induced by MMS in wild-type liver. (F) Cells were treated with MMS and 6 h later XBP1 splicing was analyzed by RT-PCR and agarose gel electrophoresis. (G) Quantification of XBP1 splicing by RT-qPCR 6 h after treatment with MMS. (H) Quantification of XBP1 splicing after treatment with MMS for the indicated time. ***p <0.001

MMS treatment led to substantial differences in gene expression between wild-type and *Aag*^−/−^ mice, with 4.7 fold more genes differentially expressed in wild-type, and with minimal overlap (Fig. 1B). This indicates that the absence of Aag affects alkylation-induced gene expression after MMS treatment.

We next performed gene-set enrichment analyses using libraries provided by the Enrichr database (25, 26). As expected from studies with cultured cells (19–21, 27), genes induced by MMS in wild-type liver are enriched for gene-sets related to ER stress (GO:0034976, p-FDR = 8.7E-06), the UPR (GO:0030968, p-FDR = 3.2E-05), and overlap with multiple gene-sets induced by drugs that are known to cause ER stress (Supplemental Data 2, Fig. S2). However, of the 12 ER stress response genes induced in wild-type, only two are also induced in the *Aag*^−/−^ liver (Supplemental Data 2). Importantly, genes induced in an Aag-dependent manner significantly overlap with genes in the transcriptional network activated by XBP1 (Fig.1C and D), a transcription factor which promotes the expression of several ER-stress related genes (22). These networks include (i) genes up-regulated in cells expressing a constitutively active form of Xbp1 (p-FDR = 3E-13; Supplemental Data 2) (28), (ii) physical Xbp1 targets according to mouse liver ChIP-seq data (p = 1.6E-14) (29), and (iii) the Xbp1 transcriptional correlation network (p = 2.5E-10) (30).

Further supporting a potential role for Xbp1 in alkylation-induced gene expression changes, genes that are downregulated following MMS treatment are enriched for independent sets of genes suppressed in cells overexpressing constitutively active Xbp1 (p-FDR = 4.1E-09 to 9.3E-05, Supplemental Data 2) (28, 31). No such enrichments are seen for genes up or down regulated in Aag-deficient livers, and Figure 1E shows that XBP1 targets are more highly induced in wild-type livers than in *Aag*^−/−^ livers, indicating that MMS induces the expression of XBP1 targets in an AAG-dependent manner.

Generation of transcriptionally active XBP1 protein requires unconventional splicing of its mRNA, a process initiated by the ER stress-induced endonuclease IRE1α (32). We asked, therefore, whether AAG might be required for maximal activation of XBP1 by MMS. Experiments were carried out with T98G cells, which are derived from glioblastoma, a type of cancer frequently treated with alkylating chemotherapy agents (33). MMS induced XBP1 splicing in wild-type T98G cells; however, when AAG expression and activity was reduced by RNAi (Fig. S3, A-C), XBP1 splicing was substantially diminished (Fig. 1F and G). Cells transfected with a non-silencing control shRNA (shNS) displayed some XBP1 splicing even in the absence of MMS, which may be due to stress caused by the CMV-driven knockdown system we employed (Fig. 1F; Fig. S4). A time curve of MMS treatment showed that XBP1 splicing peaked after 4 h, and splicing was reduced in AAG-deficient cells at all time points up to 24 h (Fig. 1H).

We also compared XBP1 splicing in A172 glioblastoma cells that express comparably low endogenous levels of AAG, versus A172 cells stably expressing a GFP-AAG fusion protein (Fig. S3). Differential AAG expression and activity in these cells was verified by qPCR, immunoblotting and enzyme assay (Fig. S3, D-F). XBP1 splicing could not be detected in parental A172 cells (Ctrl) or in A172 cells expressing just GFP (GFP), while splicing was effective in cells expressing GFP-AAG, albeit in a manner that is apparently independent of MMS treatment (Fig. S4). When *XBP1* mRNA was analyzed by qPCR, on the other hand, MMS-induced splicing was detectable in parental A172 and in GFP cells, and splicing increased about twofold in cells over expressing GFP-AAG (Fig. S4).

### AAG modulates expression of XBP1 target genes in glioblastoma cells after alkylation treatment

To verify the effects of AAG on XBP1 activation, we measured the mRNA levels of *HSPA5* (Bip/GRP78) and *HERPUD1*, two prominent markers of ER stress and known targets of XBP1 regulation (28, 29). In T98G shNS, which express abundant levels of AAG (Fig. S3), MMS treatment increases BiP and HERPUD1 mRNA 3 to 4-fold (p<0.05; p<0.01), with BiP peaking after 6 h while HERPUD1 continues to increase for up to 24 h; in *AAG* knockdown cells, by contrast, BiP is not induced by MMS, and *HERPUD1* induction is significantly reduced (Fig. 2A). Immunoblotting confirmed that MMS-induced BiP expression is lower in *AAG* knockdown cells (Fig. 2B) reaching a maximum at 6h (Fig. S5, A-B). Regulation of BiP by MMS was also studied in GFP-transfected A172 cells, which express low levels of endogenous AAG versus cells overexpressing an AAG-GFP fusion protein. Once again, AAG expression positively correlated with MMS-dependent BiP induction; BiP mRNA levels were higher in *AAG* overexpressing cells than in control cells after MMS treatment at all time points tested (Fig. S5C). Western blotting confirmed that MMS treatment induced BiP to higher levels in the AAG overexpressing A172 cells (Fig. 2C). These data further support the conclusion that MMS triggers an ER stress response through a pathway involving AAG and XBP1.

**Fig. 2.**
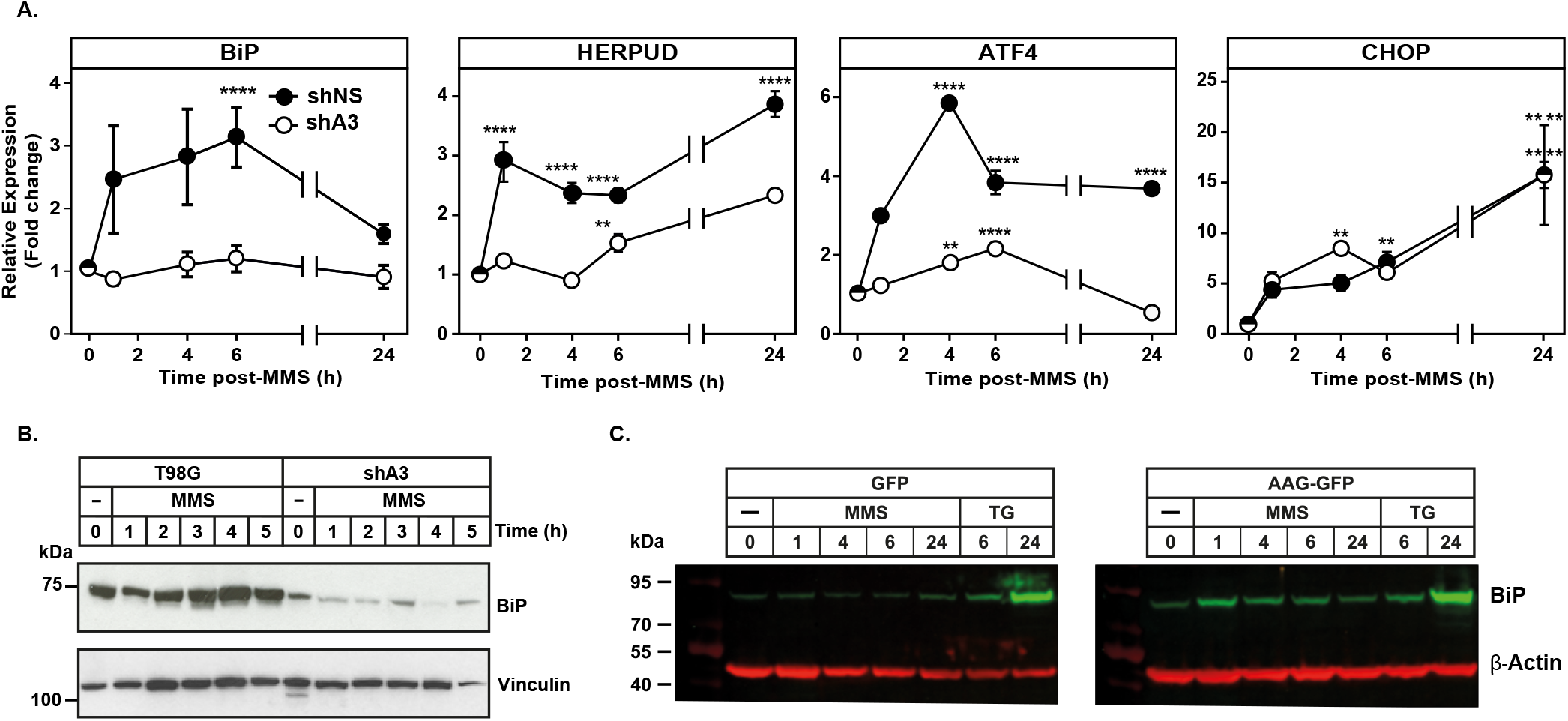
AAG modulates expression of XBP1 target genes in glioblastoma cells after alkylation treatment. Cells were treated with 2.5 mM MMS or 300 nM thapsigargin (TG), as indicated. (A) Quantification of *BiP, HERPUD, ATF4* and *CHOP* mRNA levels by qPCR. (B) BiP protein levels in shNS and shA3 cells. (C) Quantification of BiP; protein levels were normalized to β actin and expressed relative to untreated control. (D) Quantification of BiP mRNA levels in cells overexpressing control GFP or GFP-AAG. (E) Cells were treated with MMS or TG as in (D) and BiP protein levels were measured by immunoblotting. \**P* <0.05 \*\**P* <0.01, \*\*\**P* <0.001.

### The role of AAG is specific for alkylation-induced ER stress

To gauge whether AAG might be important for other branches of the UPR, we analyzed the mRNA levels of ATF4 and DDIT3/CHOP, which are controlled through the PERK and ATF6 pathways (34). Both ATF4 and CHOP were induced by MMS in control T98G cells, but only ATF4 induction was reduced in AAG-depleted cells (Fig. 2A).

Next, we studied the effects of the ER stressor thapsigargin, which depletes ER Ca^2+^ by blocking SERCA ATPases (35). Experiments with T98G and A172 glioblastoma lines expressing varying levels of AAG showed that thapsigargin induced splicing of XBP1 (Fig. S6 A), the transcription of BiP, HERPUD1, ATF4 or CHOP (Fig. S6 B-F) and increased BiP protein expression (Fig. S6 G) in an AAG-independent manner. These results are consistent with the model that AAG affects specifically alkylation-induced ER stress through a pathway that feeds into the XBP1 and likely the PERK branches of the UPR.

### Evidence for a non-canonical AAG role in alkylation-induced UPR induction

Alkylation DNA damage can affect gene expression by direct structural hindrance of the transcriptional machinery or as a consequence of DNA repair. We therefore assessed whether the DNA repair activity of AAG is required for this response. We transfected *AAG* knockdown cells with the mouse Aag (mAag) wildtype sequence (shA3_WT) or a mAag-Y147I/H156L double mutant (shA3_MUT, Fig. 3A). As control, cells were also transfected with the empty vector (shA3_EV). The mAag-Y147I/H156L double mutant is the mouse equivalent of the characterized human AAG-Y127I/H136L mutant, which is catalytically inactive despite retaining some ability to bind damaged DNA (36). We took advantage of a species mismatch that renders the shRNA sequence used for stable hAAG knockdown unable to target the mouse Aag transcript; Western Blotting analysis confirms complementation of Aag expression in these cells (Fig. 3B). As an additional measure of the ability of mAag and its mutant to repair alkylated DNA base lesions we assessed their ability to protect cells against MMS-induced cytotoxicity. Upon MMS exposure, cells expressing mAag-Y147I/H156L failed to rescue survival relative to that of cells expressing wild type mAag (Fig. 3C). To confirm complementation of glycosylase activity, we employed a fluorescently labelled oligonucleotide complex containing a single hypoxanthine as Aag substrate (labelled “Substrate”, Fig. S7). Excision of the hypoxanthine base by recombinant AAG following alkali cleavage by NaOH generates a product of the expected size (labelled “Product”). Using this assay, we can detect a time and nuclear extract concentration-dependent decrease in glycosylase activity in *AAG* knockdown cells compared to control (Fig. 3D), and this defect is complemented when cells are transfected with wild type mAag, but not with the mAag-Y147I/H156L double mutant (Fig. 3 E and F).

**Fig. 3.**
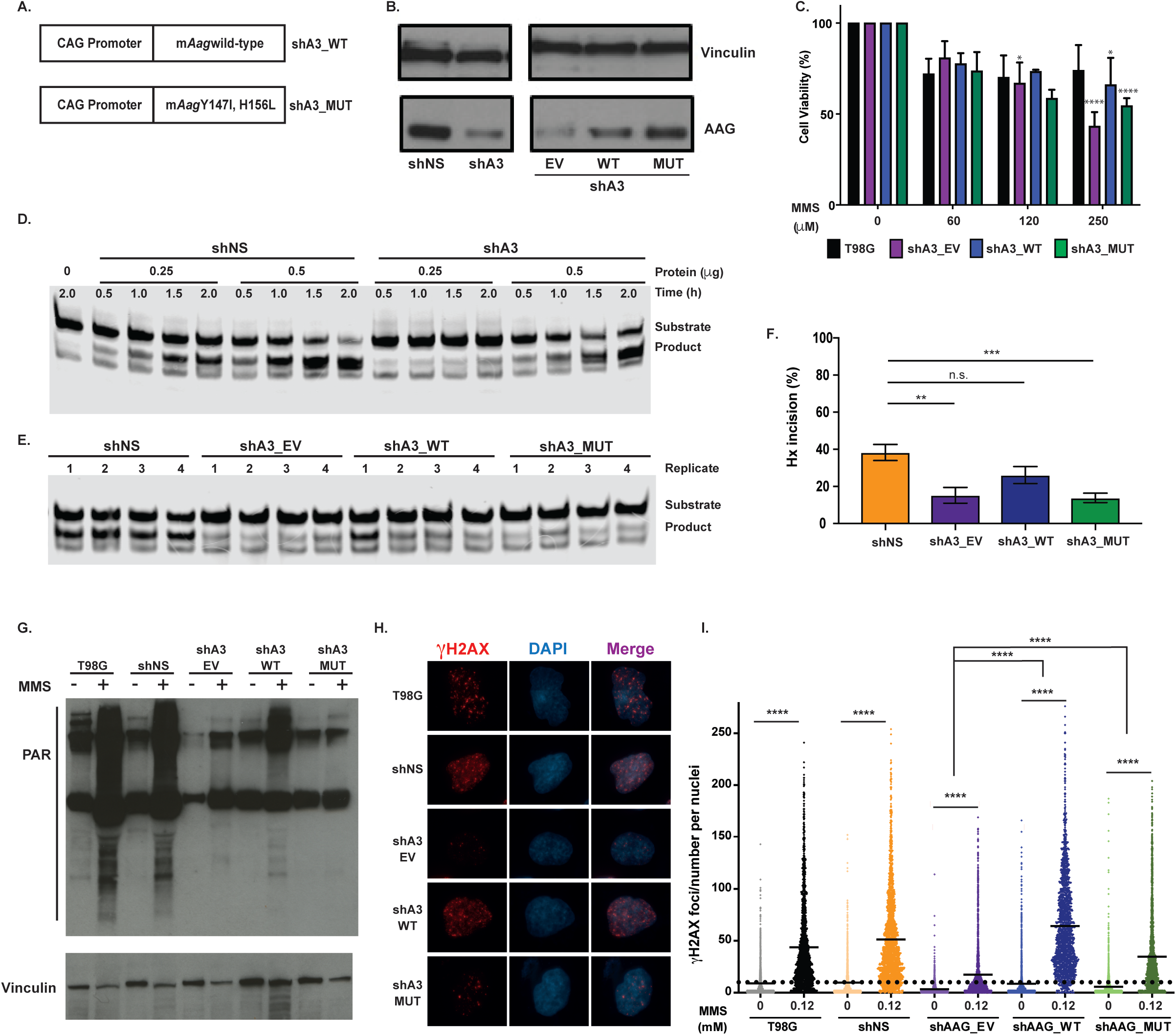
The mAag Y147I, H156L double mutant variant is defective in glycosylase activity and alkylation-induced PARylation but retains the ability to induce γH2AX foci after MMS treatment. (A) Schematic representation of constructs used for transfection of *AAG* knockdown glioblastoma cells with wild-type mouse Aag (mAag) or mAag Y147I, H156L double mutant. (B) Increased AAG expression in complemented *AAG* knockdown cells confirmed by Western blot. (C) Cell viability of shAAG cells complemented and control cells exposed to the indicated doses of MMS for 1h followed by 24h incubation in drug-free media. (D) A fluorescently-labelled Hx containing oligonucleotide (AAG substrate) was incubated with two concentrations (0.25 or 0.5 μg) of nuclear extracts for increasing times at 37 °C. Reaction products were run on a 15% denaturing polyacrylamide gel and visualized by the LYCOR Odyssey. (E) AAG activity was assessed in independent cell lysates (0.25 μg, n=4) for each genotype, after incubation with the Hx-containing oligo for 1.5 hours at 37 °C. (F) Quantification of Hx incision calculated as signal present in the product band relative to total. (G) Total PAR levels were examined by Western blotting against PAR after MMS treatment in FBS-free media for 1h followed by 1h treatment with PARG inhibitor. H - I) Representative immunofluorescence images (H) and quantification of γH2AX foci per nuclei (I) in shAAG complemented and control cells, exposed to MMS for 1h followed by 1h in drug-free media. Experiments were performed at least 3 times, *P <0.05 **P <0.01, ***P <0.001, ****P <0.0001.

AAG-initiated BER leads to the generation of repair intermediates that activate PARP (poly(ADP-ribose) polymerase) (12). PARP is activated at SSBs, to synthesize a polymer of ADP-ribose (PAR) onto itself plus acceptor proteins usually associated with DNA transactions and shaping cellular outcome to a variety of stress conditions (45, 46). Thus, we next assessed total cellular PARylation in these cell lines. Western blot analyses against PAR using total cell lysates showed that MMS treatment increased total cellular PAR levels in the control cells T98G and shNS and in the knockdown cells expressing wild type mAag, but not in knockdown cells alone or expressing the mAag-Y147I/H156L mutant (Fig. 3G).

Phosphorylation of the histone variant H2AX is a key cellular event in the DDR and induced in response to alkylation. We previously showed AAG-initiated BER is necessary for alkylation-induced γH2AX foci (14). Indeed, we confirm that *AAG* knockdown cells show reduced γH2AX foci formation after MMS treatment (Fig. 3H and I). Surprisingly, however, we observe increased levels of γH2AX foci in *AAG* knockdown cells complemented with either wild type mAag or the double mutant mAag-Y147I/H156L, albeit at reduced levels in the mutant-complemented cells (Fig. 3H and I). Together, these results suggest that despite a repair defect as shown by reduced glycosylase activity, reduced survival to alkylation and defective alkylation-induced PARylation, the mutated mAag-Y147I/H156L protein retains some ability to induce γH2AX.

Finally, we tested whether the mAag-Y147I/H156L double mutant could restore the splicing of XBP1 and expression of other markers of UPR (Fig. 4). Mutant and wildtype mAag expression restores XBP1 splicing and other markers of ER stress, at least partially (Fig. 4 A-D). Strikingly, *AAG* knock-down cells expressing the mAag-Y147I/H156L mutant display more prominent XBP1 splicing than cells complemented with wild-type mAag (Figure 4B). This suggests AAG can mediate alkylation-induced XBP1 splicing independently from its glycosylase activity and also in a PARP-independent manner. Indeed, we find that alkylation-induced XBP1 splicing is unaffected in *PARP* knockout cells (37) (Fig. S8). We propose Aag has canonical and non-canonical functions affecting cellular responses to alkylation, including ER stress induction.

**Fig. 4.**
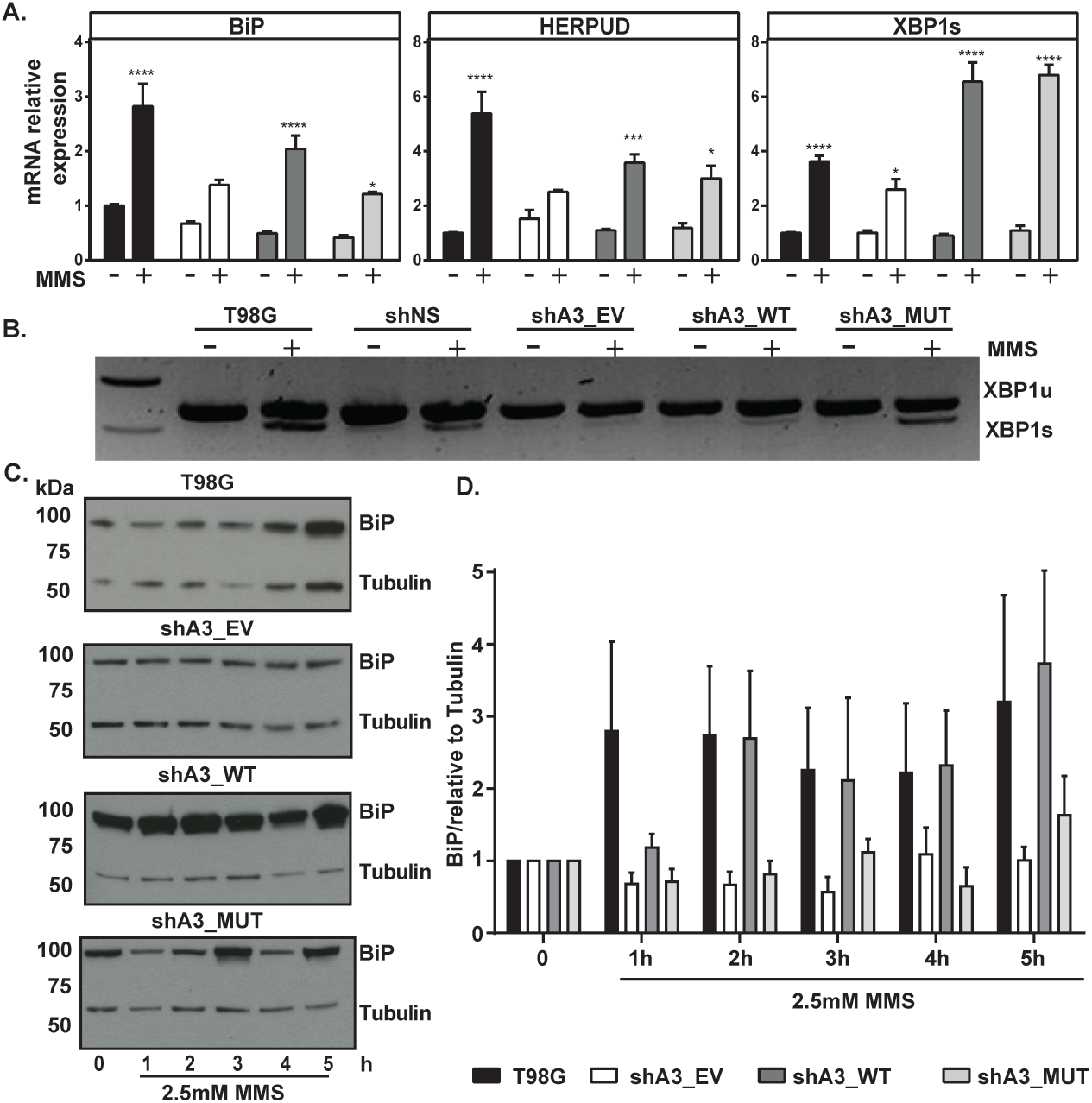
AAG glycosylase activity is not required for alkylation-induced XBP1 splicing. Cells were treated with MMS, doses as indicated. (A) Quantification of *BiP, HERPUD,* and spliced *XBP1* mRNA levels by qPCR. (B) MMS treatment induced XBP1 splicing in T98G, shNS cells and *AAG* knockdown cells complemented with wildtype and Y147I, H156L double mutant mAag. (C) Temporal characterization of BiP protein levels in complemented AAG knockdown cells and controls (D) Quantification of BiP protein, levels were normalized to α–tubulin and expressed relative to untreated control.

### AAG-mediated UPR induction plays a role in survival to alkylation

Activation of the UPR has been proposed as a mechanism underpinning glioblastoma response to treatment. UPR down regulation increases glioblastoma sensitivity to gamma radiation, etoposide and cisplatin (38–40) and ROS inducers (41). Moreover, UPR inducing drugs sensitize glioblastoma cells to the alkylating agent temozolomide (42, 43).

We therefore assessed clonogenic survival following alkylation treatment in *AAG* knockdown shA3 cells and in control T98G cells. *AAG* knockdown significantly decreased survival after treatment with MMS (Fig. 5A) or temozolomide (Fig. 5B). Importantly, cell survival was significantly reduced in AAG-depleted cells at doses of MMS and temozolomide that only modestly reduced viability in control cells. That decreased AAG levels correlate with increased alkylation sensitivity could be explained by a lower DNA repair capacity (17, 44) but contrast with multiple reports linking increased AAG levels with enhanced alkylation sensitivity (5). Nevertheless, our results showing that AAG is required for alkylation-induced UPR induction suggests alkylation sensitivity may not solely depend on DNA repair but also on the adaptive response induced by ER stress to promote survival.

**Fig. 5.**
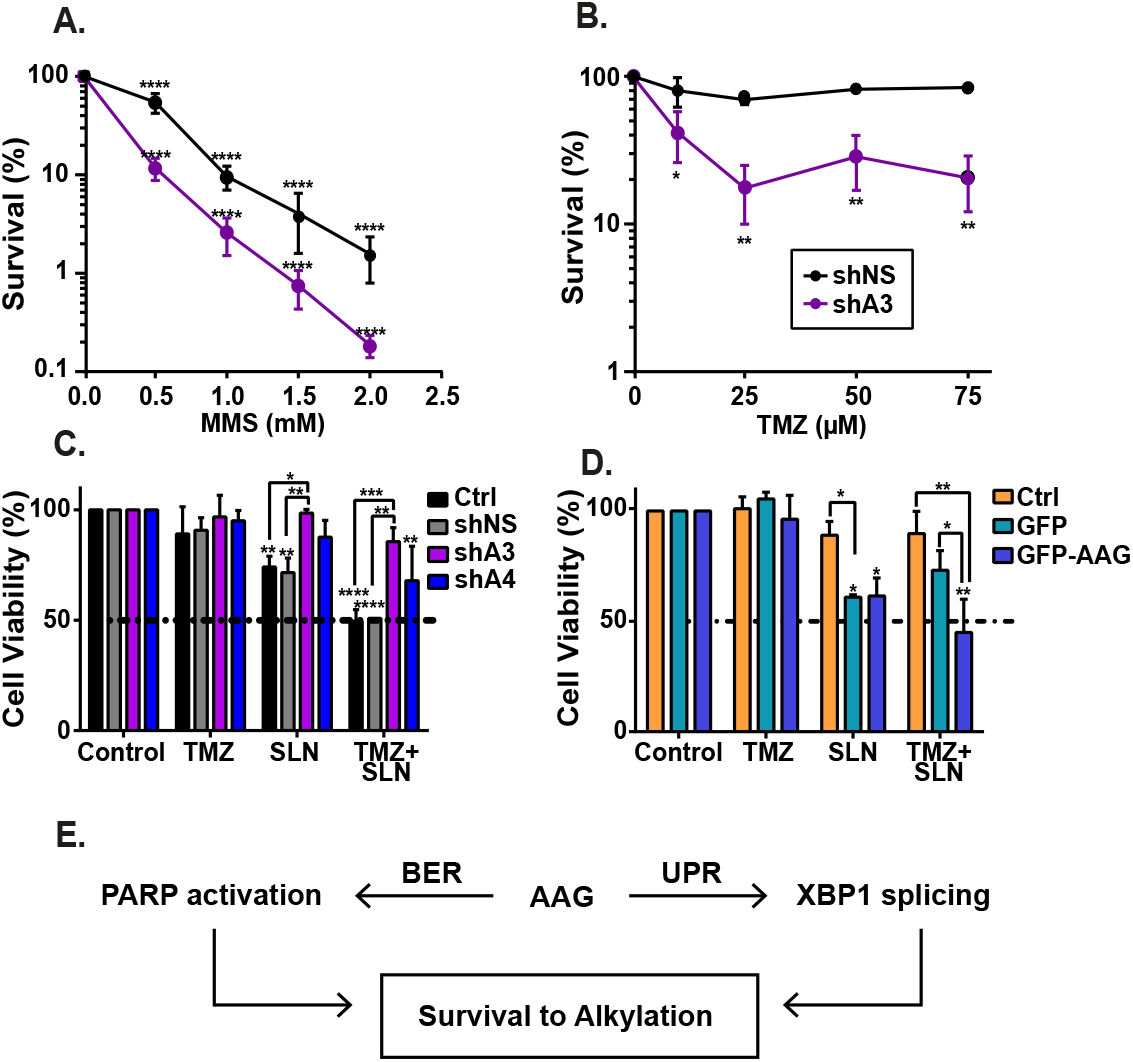
AAG-mediated UPR induction plays a role in survival to alkylation. (A and B) Clonogenic survival for cells treated with MMS (0.5 to 2 mM) for 1h or temozolomide (TMZ, 10 to 75 γM) for 5 days and incubated in drug-free media for up to 14 days. shA3 cells were more sensitive than shNS cells to (A) MMS or (B) temozolomide. (C and D) MTS survival for cells treated with temozolomide (0.2 γM) for 5 days singly or in combination with salinomycin (SLN, 0.1 γM). (C) MTS survival for shA3 and shA4 cells compared to wild type T98G (Ctrl) and shNS cells (D) MTS survival for cells overexpressing AAG (GFP-AAG) compared to cells with low endogenous AAG expression (Cntl or GFP). (E) Model for AAG’s role in survival to alkylation through independent functions in DNA repair (via PARP1 activation) and in UPR induction (via XBP1 splicing). *P <0.05 **P <0.01, ***P <0.001, ****P <0.0001.

To further address the biological relevance of alkylation-induced ER stress in this model, we next treated the glioblastoma cell lines with a non-toxic dose of temozolomide (0.2 γM), either alone or in combination with salinomycin (0.1 γM), an ionophore agent known to induce ER stress (45). Salinomycin treatment sensitizes glioblastoma cells to temozolomide, and survival after temozolomide and salinomycin co-treatment is reduced in AAG-expressing cells (Fig. 5 C and D). Strikingly, *AAG* knockdown protects cells against salinomycin, alone or in combination with temozolomide, demonstrating that AAG-mediated UPR induction contributes to cytotoxicity in this cell type. Taken together, our data are consistent with the conclusion that alkylation, signaling through AAG, induces hallmarks of an ER stress response. Whether this cascade results in cell death is likely to depend on signal strength and timing, cell type and physiological context (34).

## Discussion

Cancer chemotherapy relies on DNA damage induction by reactive compounds that are often also proteotoxic, thus increasing focus has been placed on the potential intersection between the UPR and genome damage response pathways. The present work uncovers a novel role for alkyladenine DNA glycosylase or AAG, a DNA repair enzyme, in the activation of the UPR in response to alkylating chemotherapeutic agents. We show here that alkylation treatment activates the UPR both in mouse liver and glioblastoma cells. We find that AAG modulates alkylation-induced UPR in a mechanism involving XBP1 splicing. Crucially, our results suggest this modulation does not depend on the DNA repair activity of AAG.

Alkylating agents target a variety of cellular macromolecules, including proteins. Although our study in the mouse liver examines alkylation-induced transcriptional reprogramming in repair-deficient mice, they are mirrored by studies in *Saccharomyces cerevisiae* that similarly showed transcriptional induction of the UPR by alkylation treatment (19, 46). Finally, UPR induction was shown to be important for alkylation survival in *Drosophila*, murine and human cell models (21, 27). Our work now shows that alkylation induces the UPR through a pathway that involves AAG and XBP1.

Up-regulation of UPR markers has been detected in glioblastoma and other cancer types, with implications for cancer progression and response to therapy. The IRE1α/XBP1 branch of the UPR has been implicated in glioblastoma prognosis (47), potentially by promoting glioma infiltration and motility through the modulation of pro-angiogenic and pro-inflammatory chemokines (48, 49). While our results support a role for the IRE1α/XBP1 branch in alkylation-induced UPR, we cannot rule out the participation of other UPR branches, namely PERK and ATF6. ATF6 reportedly affects glioblastoma development and radiotherapy resistance (38) while PERK is important for glioblastoma growth and survival (39). Given the importance of the UPR in cancer, a better understanding of how AAG affects alkylation-induced UPR could advance efforts for therapeutically targeting ER stress/UPR in cancer.

Our gene expression analyses in the MMS-treated mouse liver show that the transcriptional response to alkylation treatment is profoundly reduced in the absence of Aag. Besides the enrichment for ER stress/UPR related transcripts, and the overlap with multiple gene-sets induced by ER-stress inducing drugs (Fig. S2), we find that genes induced by MMS in wild-type liver also significantly overlap with single gene perturbations associated with specific biological phenotypes related to ER redox homeostasis (overexpression of *ERO1L*), UPR (*H6pd* knock-out) and recovery from inflammation and toxicity (*Socs3*, *Mat1a* and *Txnrd1* knock-outs) (Supplemental Data 2). In contrast, Aag-deficient mice do not exhibit alterations in the expression of these markers of tissue injury. This is consistent with previous work showing that *Aag* knockout protects from alkylation-induced cell death (16), and with a role for Aag in promoting alkylation-induced tissue injury.

AAG-mediated alkylation-induced toxicity is rescued in animals that lack Parp-1 (16). It is thought that AAG-initiated BER followed by DNA strand cleavage induces PARP-1 activation that results in tissue damage by depleting cells of energy, which leads to necrosis. It is not clear whether AAG-mediated UPR is an additional pathway of cell death induction that contributes to alkylation-induced tissue damage. Nevertheless, our results suggest PARP-1 activation is not necessary for AAG-mediated UPR, at least as it relates to XBP1 splicing. It is worthy of mention that PARP-1 has been previously shown to promote enhanced activity of the 20S proteasome in response to oxidative damage, thus contributing to removal of oxidised nuclear proteins (50, 51). Our results do not exclude the possibility that PARP-1 could play a similar BER-independent role in response to alkylated proteins.

Our data indicate that a mutant AAG variant defective in base excision nevertheless complements some alkylation-responsive phenotypes as measured in AAG-deficient cells, namely γH2AX foci formation and XBP1 splicing. This indicates that alkylation-induced γH2AX foci formation may occur, at least in part, independently of AAG base excision activity, and that the connection between the UPR and AAG can proceed in the absence of BER-initiation and PARP-1 activation. This is an important consideration because increased AAG levels have been associated with increased inflammation and increased genetic instability in the form of increased microsatellite instability (MSI) (52). Moreover, overexpression of the corresponding human AAG-Y127I/H136L double mutant led to increased MSI in cultured human K562 cells (36). Our results showing that a catalytically inactive AAG protein can still induce γH2AX and the DDR suggest that some of the effects of AAG may derive from its ability to recognise damaged bases. Recently, a catalytically inactive mutant of 8-oxoguanine DNA glycosylase (OGG1) was shown to act as a potent regulator of gene expression, and substrate binding was required for OGG1-driven transcriptional activation (53). Similarly, AAG-mediated recognition of alkylated bases could initiate signaling that propagates towards XBP1 splicing. Alternatively, AAG may have a role in alkylation-induced UPR that is independent of DNA binding. Our results are consistent with a model where AAG plays independent roles in BER and UPR induction (Fig. 5E), but further studies are needed to establish whether substrate binding is required for alkylation-induced UPR activation.

The biochemical properties of the AAG protein of binding to and excising damaged bases have important implications for the dynamics of DNA transactions (e.g. repair, replication and transcription) in the presence of physiological or supra-physiological levels of DNA damage. Our results support a model where AAG exerts its effects through binding damaged DNA and interacting with other proteins associated with DNA transactions. It has been reported that wild type AAG interacts with a number of other proteins with known roles in transcriptional modulation and/or ubiquitin mediated proteolytic pathways, including HR23A, HR23B, p53 and estrogen receptor alpha (ERα) (54–56). More recently, AAG was shown to form a complex with active RNA pol II through direct binding to the ELP1 subunit of the transcriptional elongator complex (57). AAG is also reported to interact with MBD1, a methyl-CpG binding domain protein implicated in transcriptional regulation (58), and ubiquitin-like with PHD and RING Finger domains 1 and 2 (UHRF1/2) proteins, “hub” proteins involved in epigenetic regulation (59). Therefore, AAG could affect the UPR through its interaction with proteins important for ER stress response or transcriptional control in general.

DNA repair and the UPR were previously reported to cooperate in response to cellular stress. In particular, an important subset of XBP1 targets are DNA repair genes (60). ER stress induction was reported to modulate expression of BER proteins, including AAG (61, 62) and APEX1 (63). Further, pharmacological ER stress induction potentiated the cytotoxic effects of temozolomide in glioblastoma cells (61, 62). These results implicate ER stress/UPR in DNA repair modulation and indicate the two pathways cooperate in stress response.

Together, our results demonstrate the DNA repair enzyme AAG plays a role in alkylation-induced UPR activation. Whether there is a direct effect of AAG on alkylation-induced UPR or whether it is due to AAG-mediated DNA damage recognition remains to be defined, but our data strongly suggests AAG’s base excision activity is not required. We anticipate that a detailed mechanistic dissection of this stress response crosstalk will lead to a better understanding of cellular outcomes upon alkylation exposure and may shape future advances in the prevention and treatment of cancer and other age-related diseases.

## Materials and methods

### Materials

MMS, puromycin, salinomycin, temozolomide and DMEM low glucose were from Sigma (St. Louis, MO). Cell culture reagents were from Invitrogen (Carlsbad, CA).

### Mice

*Aag* null mice were described previously (24). Details about animal experiments are described in *Supporting Methods.* The MIT Committee for Animal Care (CAC) approved all animal procedures.

### Microarray processing and data analysis

Messenger RNA was isolated, amplified and hybridized onto Affymetrix GeneChip Mouse Genome 430A 2.0 arrays according to the protocols suggested by Affymetrix (Santa Clara, CA). Data were analysed using R/Bioconductor as described in *Supporting Methods*.

### Plasmids

Plasmid pEGFP-hAAG was generated by cloning the XhoI flanked *AAG* cDNA from pCAGGS-hAAG (36), into pEGFP-C3 (Clontech, Takara BioUSA, Inc). Plasmid pCAGGS with the mAag-Y147I/H156L double mutant cDNA was generated by site-directed mutagenesis of the wild-type cDNA, as previously described (36). The nucleotide sequence of all plasmids was confirmed by DNA sequencing. Lentiviral shRNA plasmids based on pGIPZ were purchased from Dharmacon; insert sequences are listed in Supplemental Table S1.

### Cell culture

A172 and T98G human glioblastoma cell lines were obtained from ATCC, were free from mycoplasma contamination and always used from a young stock. Experimental details related to cell culture and cell line development and complementation are described in *Supporting Methods*.

### AAG Activity assay

AAG activity assay is described in *Supporting Methods*.

### XBP1 Splicing and expression analysis

Quantitative and qualitative PCR methods and reagents are described in *Supporting Methods.*

### Immunoblotting, PAR andγH2AX detection

For details on immunoblotting and antibodies, see *Supporting Methods*.

### Statistical analysis

Unless otherwise stated, results are presented as mean ± standard error of the mean and analyses are results of three or more independent experiments. Statistical calculations were performed with GraphPad Prism software, Version 6.0.

## Supporting information

Gene Expression Data

Enrichment Analysis

## Acknowledgments

We thank Rebecca Fry, Katherine Pease, Jennifer Calvo, Fugen Li and Stuart Levine (MIT BioMicroCenter) for help with the gene expression analysis, and Joanna Klapacz for the plasmid containing the double mutant Aag variant. We thank Miriana Machado (UFRGS, Brazil) for clerical support. The National Institutes of Environmental Health Sciences (ES11399) funded work performed at MIT. Work otherwise supported by the Royal Society (RC3532) and the Conselho Nacional de Desenvolvimento Científico e Tecnológico (CNPq, 400557/2013-4 and 207503/2015-0). LM, CFC and RA are recipients of SwB studentships (CNPq). LBM was supported by an international visiting professorship grant (CNPq). The authors declare no conflict of interest.

