## Supplemental Text and Figures for "A DNA repair-independent role for alkyladenine DNA glycosylase in alkylation-induced unfolded protein response"

**This PDF file includes:**

Supplementary text
Figs. S1 to S8
Tables S1 to S2

**Other supplementary materials for this manuscript include the following:**

Supplemental Data 1 – Detailed gene expression data
Supplemental Data 2 – Gene enrichment analysis

**Supplementary Information Text**

**Material and Methods**

**Animal Experiments.** Wild type and *Aag* null 6-8 week old male mice were used (n=3 for microarray experiments). Mice were fed standard diet *ad libitum* and housed in accredited facilities. Euthanasia was by $CO_2$ asphyxiation.

**MMS treatment of mice.** MMS was dissolved in 10% ethanol in phosphate-buffered saline (PBS). Animals were injected intraperitoneally (i.p.) with solvent or a single dose of 75 mg MMS per kg body weight, a dose known to be sub-lethal to both *Aag*-null and wild-type animals (1). Whole liver (mixtures of all 5 different liver lobes) was collected 6 h after treatment, cut into pieces, flash frozen in liquid nitrogen and stored at -80ºC until further processing.

**Microarray processing and data analysis.** Raw data background correction (RMA method), normalization (qspline method), probe specific correction, and summary value computation were performed using the affy::expresso function in R [PMID: 14960456]. Differential gene expression was studied using limma [PMID: 25605792], applying a minimum log2 fold change of 1.75 and a maximum adjusted p value (false discovery rate [FDR] method) of 0.05. Lists of differentially expressed genes were analysed for enrichment (hypergeometric test) with prior knowledge categories using 89 gene-set libraries from the enrichr website [PMID:23586463] [ PMID:27141961]; p values were corrected for multiple hypothesis testing using the FDR method. Microarray data have been deposited on GEO (GSE115254). R scripts and additional data files used in this study are available on figshare [https://doi.org/10.24376/rd.sgul.6429092] [Link while manuscript is under review: https://figshare.com/s/8ab1e67261965796151b].

**Cell culture and cell line construction.** Medium A refers to DMEM low glucose containing 10% foetal bovine serum, 100 U/mL penicillin, 100 μg/mL streptomycin, and 2 mM L-glutamine. Cells were grown in medium A at 37°C in 5% CO2. Lentivirus expressing AAG shRNAs were produced as described previously (2), followed by infection of T98G cells and selection in medium A plus 1 μg/mL puromycin. A172/hAAG cells, stably expressing EGFP-hAAG, were generated by Lipofectamine LTX (Invitrogen) mediated transfection of A172 cells with pEGFP-hAAG, followed by selection in medium A plus 600 μg/ml G418. Puromycin resistant *AAG* knockdown shAAG3 cells were complemented by transfection with the pCAGGS empty vector or the indicated construct, as well as the empty pEGFP-C3 for G418 resistance, using Lipofectamine 2000 (Invitrogen), at a ratio of 10:1 (10 parts gene expression plasmid: 1 part antibiotic selection marker plasmid). Post-transfection, cells were selected with 1000 μg/ml G418. Selected cells were cultured in the presence of 0.5 μg/mL puromycin and 400 μg/ml G418 for maintenance.

**Cell treatment and survival determination.** Clonogenic survival assays were performed as described (3). Briefly, cells were plated and treated with MMS (0.5-2mM) for 1 h or temozolomide (10 to 75μM) for 5 days. After 14 days, colonies were fixed with methanol and stained with 0.1% crystal violet. Colonies containing >50 cells were scored and the percent survival calculated relative to untreated control. For viability, cells were pre-treated with salinomycin (0.1 μM) for 24 hours and then temozolomide treated in the presence of salinomycin for 5 days. Cell viability was determined after treatment using the MTS-based CellTiter 96® AQueous One Solution Cell Proliferation Assay (Promega, #G3580).

**MMS sensitivity assay for complemented cells**

Sensitivity assay for complemented *AAG* knockdown cells and controls was performed as described(4). Briefly, cells were seeded in triplicate in a clear bottom black 96-well microplate at a density of 5000 cells per well. MMS treatment was for 1h in serum-free media at concentrations ranging from 0 (DMSO) to 0.25 mM. Post-treatment, media was replaced by complete media and cells were incubated for 24h. Images of entire wells were acquired at 4 x magnification with a Cytation 5 Cell Imaging Multi-Mode Reader followed by quantification of Hoechst-stained nuclei with the Gen5 Data Analysis Software v3.03 (BioTek Instruments). Cell viability was expressed as percentage of survival in MMS-treated cells relative to vehicle (DMSO)-treated cells. Results represent the mean ± SEM of at least 3 independent experiments.

**AAG Activity assay**. Briefly, nuclear extracts were prepared from cells for AAG activity analysis using a NE-PER™ Nuclear and Cytoplasmic Extraction Reagents kit and the concentration of protein in the extracts determined using the MicroBCA kit according to the manufacturer's instructions (ThermoFisher Scientific, Loughborough, UK). Synthetic oligonucleotides HX02 (sequence 5' Phos-CACGAITCAACTCAGCAACTCC*T*T-$NH_2$ 3', where Phos indicates a phosphate modification, I indicates an internal inosine, * indicates a phosphorothioate linkage between nucleotides and $NH_2$ indicates an amino group modification) and Loop01Aird (5' IRDye-T*T*GGAGTTGCTGAGTTGATTCGTGAGCACCAACCGGTGCT 3', where IRDye indicates a IRDye® 800CW modification) were synthesized and HPLC purified by Integrated DNA Technologies (Leuven, Belgium). The plate based glycosylase assays were done as previously published (5). For assessing glycosylase activity of knockdown cells complemented with wildtype or mutant mouse *Aag*, a gel-based assay was used. Briefly, 3 μM HX02 was annealed to 3 μM Loop01AIRD in a buffer containing 30 mM Tris HCl and 10 mM $MgCl_2$ with total reaction volume 40 μl. Annealing was done by using a thermal cycler programmed as 95°C for 10 min; cool to 80°C at 1°C/min; 80°C for 10 min; cool to 70°C at 1°C/min; 70°C for 10 min; cool to 21°C at 1°C /min. After annealing, 10 mM DTT, 1 mM ATP, and 6 U T4 DNA ligase (Promega, Southampton, UK) were added to the reaction mixture and incubated for 1 h at 30°C, and then overnight at 4°C. The ligated oligo (substrate) was ethanol precipitated using 0.3 M sodium acetate and 1 μg/μl glycogen (Ambion), washed in 70% ethanol and resuspended in ultrapure water (40 μl). For the glycosylase assay reactions, the oligo substrate (3 pmol) was incubated at 37°C for times from 30 minutes to 2 h (in 30 min increments) with either recombinant hAAG (New England Biolabs, Hitchin, UK) or with nuclear extracts diluted in PIPES buffer (50mM PIPES (pH 6.7), 30mM KCl, 2mM EDTA, 3.4% v/v glycerol) in a 10 μl reaction. Following the repair incubation, the substrate was denatured by incubation with 0.1M NaOH for 20 min at 70°C, an equal volume of 2 X formamide loading buffer was added (95% formamide, 5 mM EDTA, 0.1% OrangeG) and samples incubated at 65°C for 5 min before gel electrophoresis. Reaction products were resolved in a 15% polyacrylamide/7M urea gel electrophoresed in TBE buffer. Gels were imaged using the Odyssey CLx IR imaging system (LI-COR Biosciences, Lincoln, USA), and image acquisition and signal quantification were done using Image Studio software (LI-COR Biosciences, Lincoln, USA).

**XBP1 Splicing** The detection of XBP1u (unspliced) and XBP1s (spliced) transcripts was performed as described in (6), except that Go Taq Green (Promega) was used, and PCR products were resolved on a 2.5% agarose gel.

**mRNA Expression Analysis** RNA was extracted using the PureLink RNA Mini kit (Invitrogen, Carlsbad, USA) and first strand cDNA was synthesized using the Maxima First Strand cDNA Synthesis (Thermo Fisher Scientific, Waltham, USA), according to the manufacturer's instructions. RNA amplification was done using SYBR Green Maxima SYBR Green/ROX qPCR Master Mix (Thermo Fisher Scientific, Waltham, USA) and quantitative real-time PCR was performed using QuantStudio 7 Flex Real-Time PCR System (Life Technologies, Carlsbad, USA). All experiments were performed with biological and technical triplicates. Results were generated using the comparative Ct method and are expressed as fold change relative to the untreated control (7). A complete list of primers used in this study is shown in Table S2.

**Immunoblotting** Cells were lysed with M-PER Mammalian Protein Extraction Reagent (Thermo Fisher Scientific, Waltham, USA), supplemented with 1X Phosphatase inhibitor (Sigma-Aldrich, St. Louis, USA) and 1x protease inhibitor cocktail (Thermo Fisher Scientific, Waltham, USA). Protein concentration was determined using the BCA assay (Pierce). Total protein lysates (20 μg) were separated under denaturing conditions in 4-20% True-PAGE (Sigma-Aldrich) gels. Proteins were then transferred onto PVDF membranes (Life Technologies). Membranes were blocked in 1% nonfat milk and incubated overnight with specific antibodies at 4 °C. Antibodies were as follows: β-actin (1:10,000 dilution; ab52614, Abcam), AAG (1:500 dilution; HPA006531, Sigma-Aldrich), BiP (1:1,000 dilution; 3183S, Cell Signaling). After primary antibody incubation, membranes were washed and then incubated with the secondary antibodies IRDye 680RD green goat anti-rabbit IgG and IRDye 800CW red goat anti mouse IgM (LI-COR Biosciences, Lincoln, USA) at 1:10,000, for 1 hour at room temperature. Proteins were detected using the Odyssey CLx IR imaging system (LI-COR Biosciences, Lincoln, USA).

**PAR detection** Cells (1 x 10^6 cells/well) were seeded in 6-well plates and treated for 1h with MMS (1 mM) in serum-free media. After treatment, media was replaced with complete media containing 1 μM PDDX004 (N02214) (PARG inhibitor) for 1h. Cell lysates were obtained using Laemmli buffer (3 X: 150 mM Tris-HCl pH 6.8, 6% SDS, 0.3 % Bromophenol Blue, 30% Glycerol). Proteins were separated by SDS–polyacrylamide gel electrophoresis using pre-cast gels 4-12% (Invitrogen) and transferred to Nitrocellulose membrane. PAR detection was performed using a rabbit anti-poly(ADP-ribose) LP-9610 in a 1:5000 dilution (8). Mouse monoclonal anti-vinculin (1:200 000; Sigma, #V9131) was used as loading control. Horseradish peroxidase-conjugated anti-rabbit or anti mouse IgG (1:10000; Jackson Immuno Research) were used as secondary antibodies. Signal was detected using enhanced chemoluminescence (Perkin-Elmer).

**γH2AX Foci Quantification** Cells were seeded into Corning 96-Well Half Area High Content Imaging Film Bottom Microplate at 10 000 cells/well. After 24h, cells were treated with MMS in serum-free media for 1h. Cells were then incubated in drug- free complete media for 1 h. Unless otherwise stated, all immunofluorescence dilutions were prepared in PBS and incubations performed at room temperature with intervening washes in PBS. Cell fixation was carried out by incubation with 4% paraformaldehyde for 10 min followed by 100% ice-cold methanol for 5 min at −20◦C. Cells were permeabilised in 0.2% Triton X-100 for 5 min followed by a quenching step using 0.1% sodium borohydride for 5 min. After blocking for 1 h in a solution containing 10% goat serum and 1% BSA, cells were incubated for 1 h with primary antibody anti-phospho-Histone H2A.X (Ser139) (1:5000, Millipore, #05-636) diluted in 1% BSA. Secondary antibody labelling used Alexa Fluor 568 goat anti-mouse (Invitrogen, #A-11004) diluted at 1:1000 in 1% BSA for 1 h. Nuclei were stained for 10 min with 1 mg/ml 4,6-diamidino- 2-phenylindole (DAPI). Z-stack images were acquired on a ZEISS Celldiscoverer 7 automated microscope using a 50× water immersion objective, and analysed for γH2AX foci formation with ZEN Blue software 3.2 (ZEISS).

**Table S1. shRNA sequences used for silencing.**

| |
|---|
| V3LHS_343111: ACAGCTTCATCCTGTGCCA |
| V3LHS_343113: TCATGCAGAAGTACATGCC |
| V3LHS_343114: CTAGCTGGTCGCTGCTTCT |
| V3LHS_343116: CATGCAGAAGTACATGCCG |

**Table S2. RT-qPCR primers.**

| | |
|---|---|
| *ACTB* (NM_001101) | Fwd 5’ ATT GCC GAC AGG ATG CAG AA 3’<br>Rev 5’ GCT GAT CCA CAT CTG CTG GAA 3’ |
| *AAG* (NM_001015054*)* | Fwd 5’ CCC CGC AAC CGA GGC ATG TT 3’<br>Rev 5’ AGC AAG ACG CAA GCC CCG TC 3’ |
| *ATF4* (NM_001675) | Fwd 5’ TTCTCCAGCGACAAGGCTAAGG 3’<br>Rev 5’ CTCCAACATCCAATCTGTCCCG 3’ |
| *HSPA5* (NM_005347) | Fwd 5’ CTGTCCAGGCTGGTGTGCTCT 3’<br>Rev 5’ CTTGGTAGGCACCACTGTGTTC 3’ |
| *XBP1*s (NM_005080) | 412–431 5’ CCTTGTAGTTGAGAACCAGG 3’<br>834–853 5’ GGGGCTTGGTATATATGTGG 3’ |
| *HERPUD1* (NM_001010989) | Fwd 5’ CCAATGTCTCAGGGACTTGCTTC 3’<br>Rev 5’ CGATTAGAACCAGCAGGCTCCT 3’ |
| *DDIT3* (NM_004083) | Fwd 5’ GGTATGAGGACCTGCAAGAGGT 3’<br>Rev 5’ CTTGTGACCTCTGCTGGTTCTG 3’ |
| *Actb* (NM_007393) | Fwd 5’CATTGCTGACAGGATGCAGAAGG<br>Rev 5’ TGCTGGAAGGTGGACAGTGAGG |
| *Herpud1* (NM_022331) | Fwd 5’ CCTCCAAAATGCCAGAAACCAGC<br>Rev 5’ GCCGTAAACCATCACTTGAGGAG |
| *Ddit3* (NM_007837) | Fwd 5’ GGAGGTCCTGTCCTCAGATGAA<br>Rev 5’ GCTCCTCTGTCAGCCAAGCTAG |

#### Supplemental Figure S1

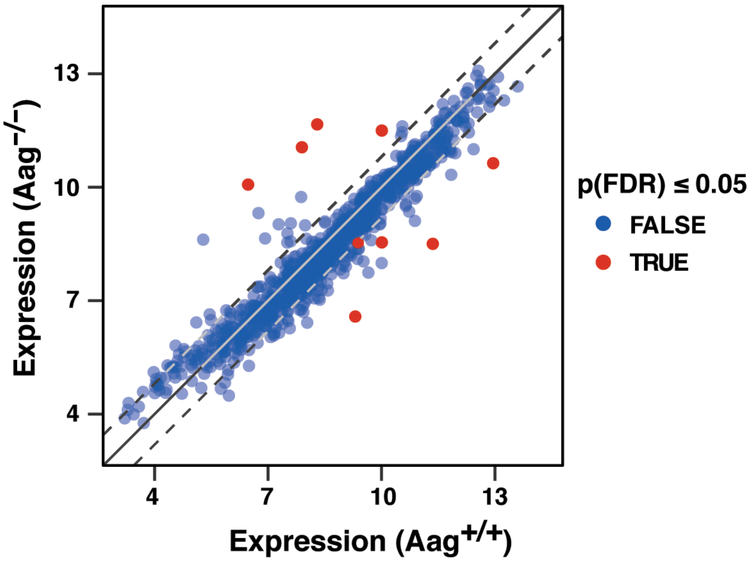


**Figure S1. Similar levels of gene expression in wild-type and Aag-deficient liver under control conditions.** Liver transcriptomes were studied by microarray analysis as described for Fig. 1. Average expression values from control-treated knockout animals are plotted against expression values from control-treated wild-type animals. Each marker represents one of 980 microarray probes whose expression changed significantly according to Figure 1 A and B, under any of the conditions. Markers are colour-coded to indicate whether the difference in expression between knockout and wild-type samples was associated with an FDR-adjusted p-value of 0.05 or less. Solid line, bisection; dashed lines, bisection plus or minus $\log_2(1.75)$.

#### Supplemental Figure S2

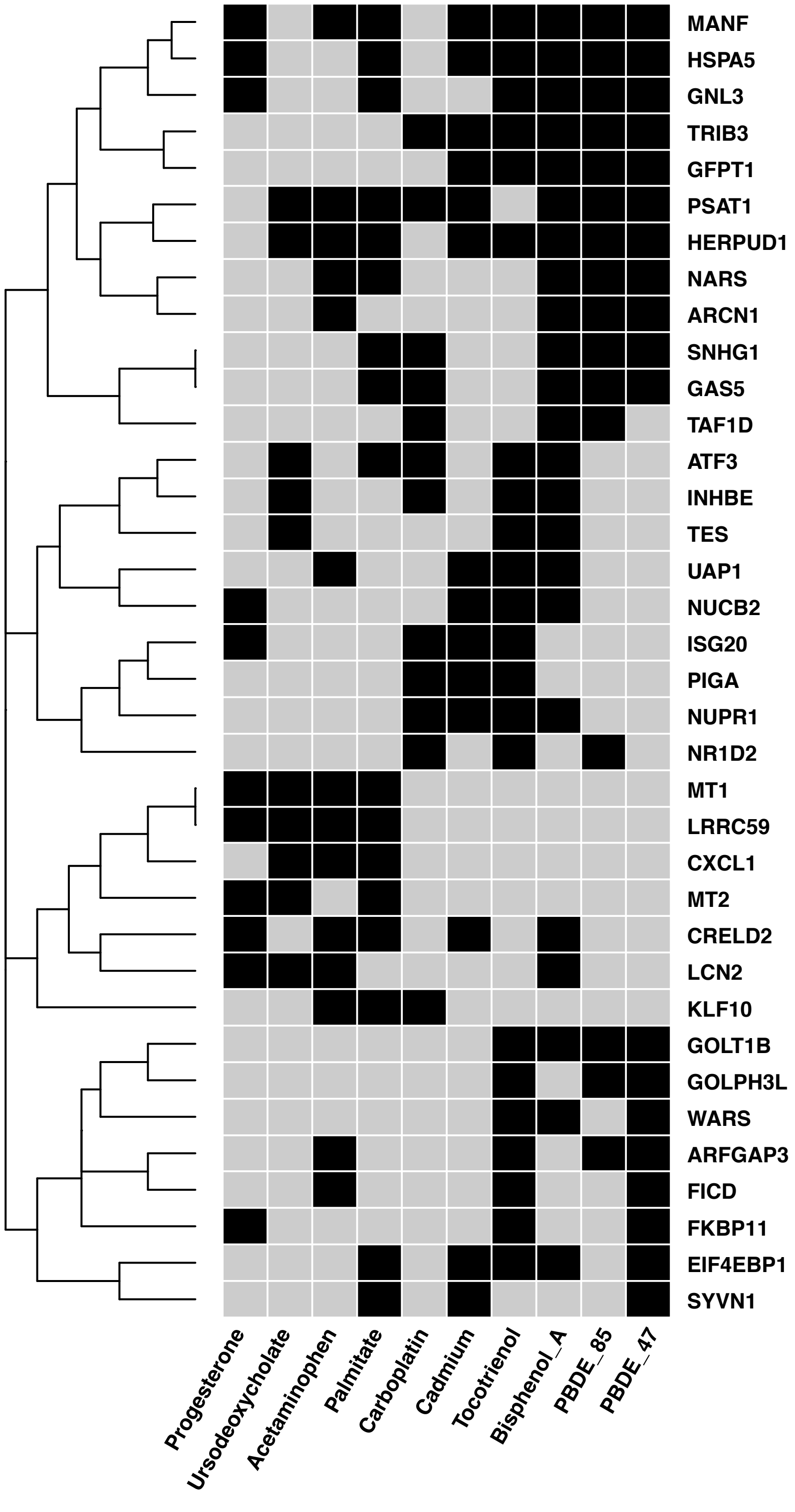


**Fig. S2**. **Genes induced by MMS and ER stress**. Genes induced by MMS in wild-type liver showed significant overlap with multiple genesets induced by ER stress-inducing compounds (Suppl. Data 2). In this binary heatmap the black squares indicate genes (y-axis) that are both induced by MMS in wild-type liver as well as by the respective ER stress-causing compound (x-axis). Only a subset of 36 genes are shown that overlapped with at least three genesets. For the complete results and methods see supporting documentation at https://github.com/nohturfft/SuppMat_Milano_et_al_2021_Biorxiv.

### Supplemental Figure 3

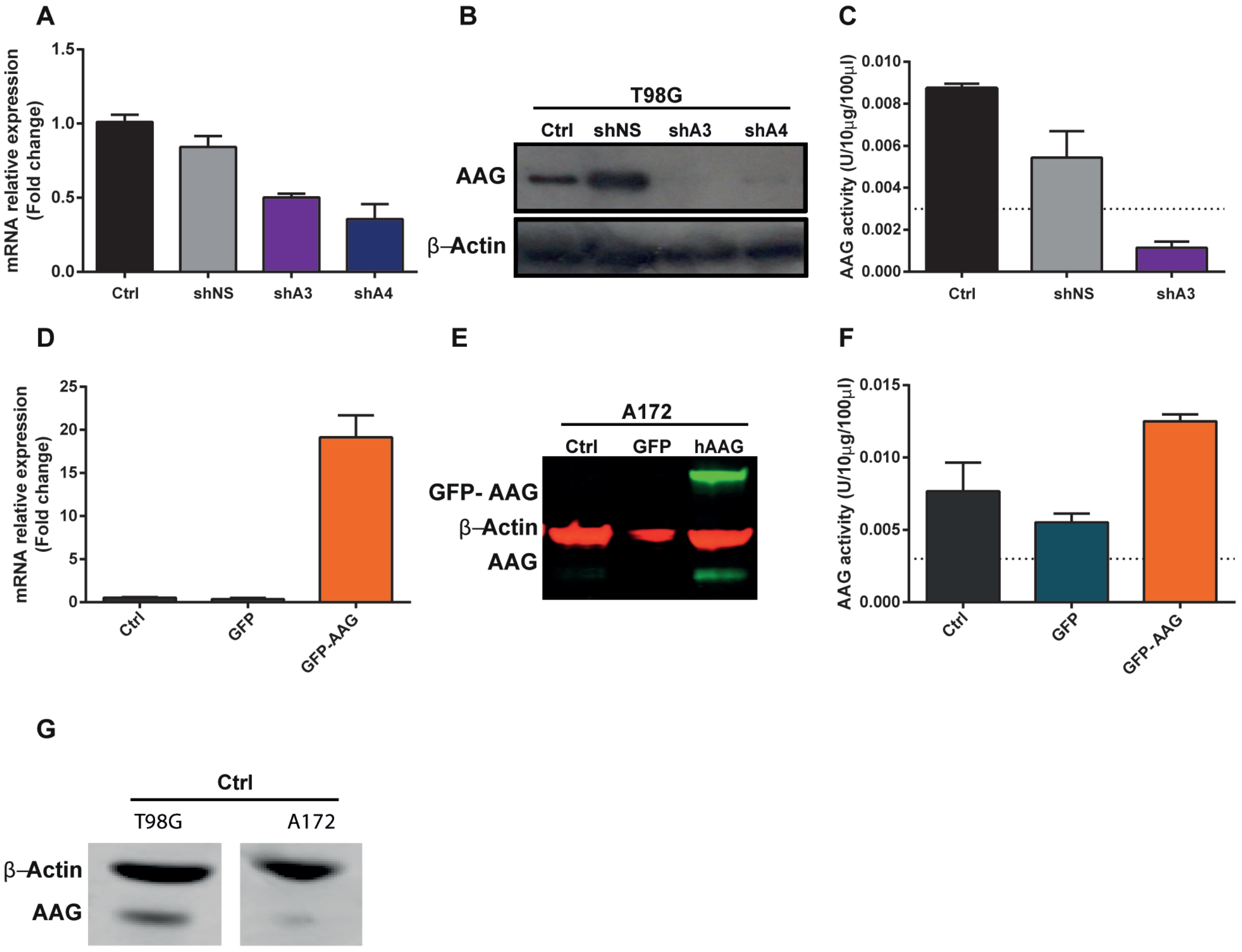


**Fig. S3. Characterization of *AAG* levels in glioblastoma cells.** T98G cells were transduced with viral particles produced from pGIPZ Lentiviral shRNA system for AAG knock down and for overexpression A172 cells were transfected with either pEGFP-C3 expressing the human *AAG* gene or empty pEGFP-C3 plasmids. (A and D) mRNA levels of AAG measured by qPCR in (A) T98G and (D) A172 cells. (B and E) Representative immunoblot of AAG levels in parental (B) T98G (WT), cells expressing scrambled shRNA control (shNS – non silencing) and cells expressing specific shRNA for AAG (shA) and (E) parental A172 (WT), pEGFP-C3 empty vector (GFP) and stable overexpressing human *AAG* (GFP-AAG), for both β actin was used as loading control. (C and F) AAG enzymatic activity in (C) T98G and (F) A172 cells. (G) Comparison of AAG protein levels in parental A172 and T98G cell lines. Results indicate the mean ± S.E. of three independent experiments.

### Supplemental Figure 4

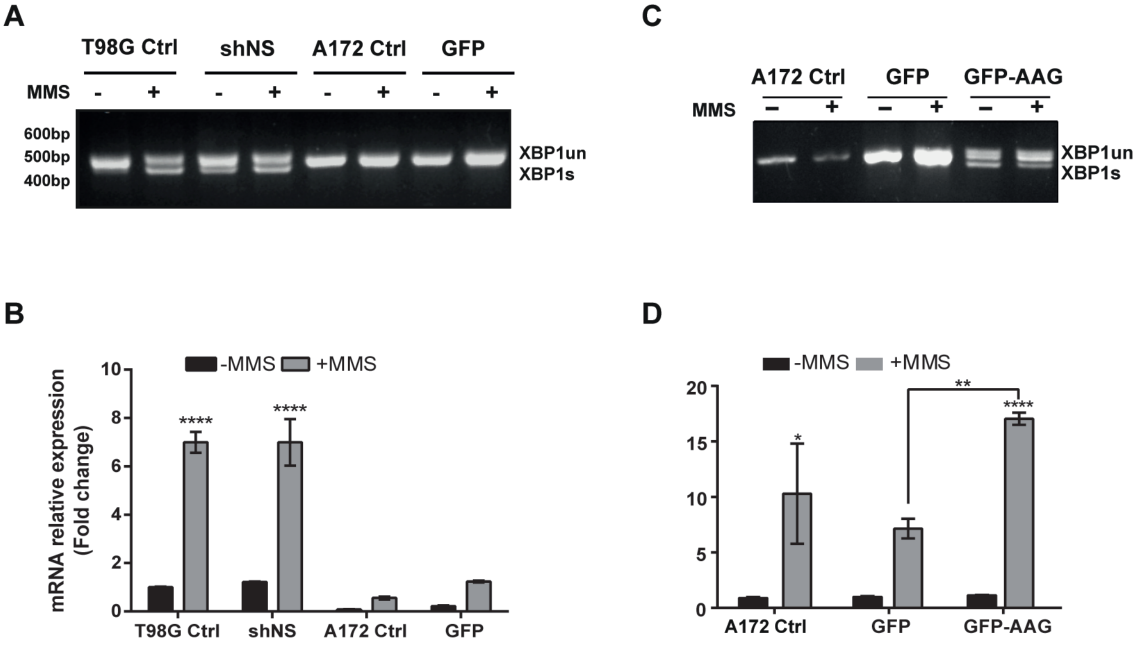


**Fig. S4 - MMS treatment induces Xbp1 splicing in cells expressing AAG.**
In A and B, comparison of MMS inducing XBP1s in T98G and A172 parental cell lines and controls; in C and D, comparison of MMS inducing XBP1 splicing in the A172 cell panel (parental, overexpressing GFP or GFP-AAG) by (A and C) RT-PCR and (B and D) qPCR. ****p<0.0001

### Supplemental Figure 5

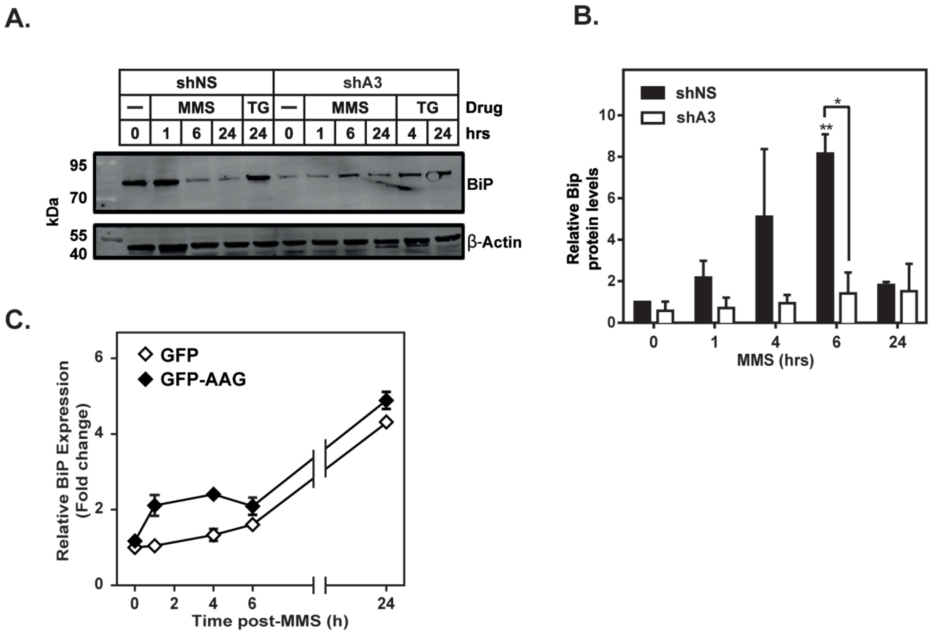


**Fig. S5.** (A) BiP protein levels in T98G and shA3 cells; (B) Quantification of BiP; protein levels were normalised to β-actin and expressed relative to untreated control; (C) Quantification of BiP mRNA levels in AAG overexpressing GFP-AAG and control GFP cells.

#### Supplemental Figure 6

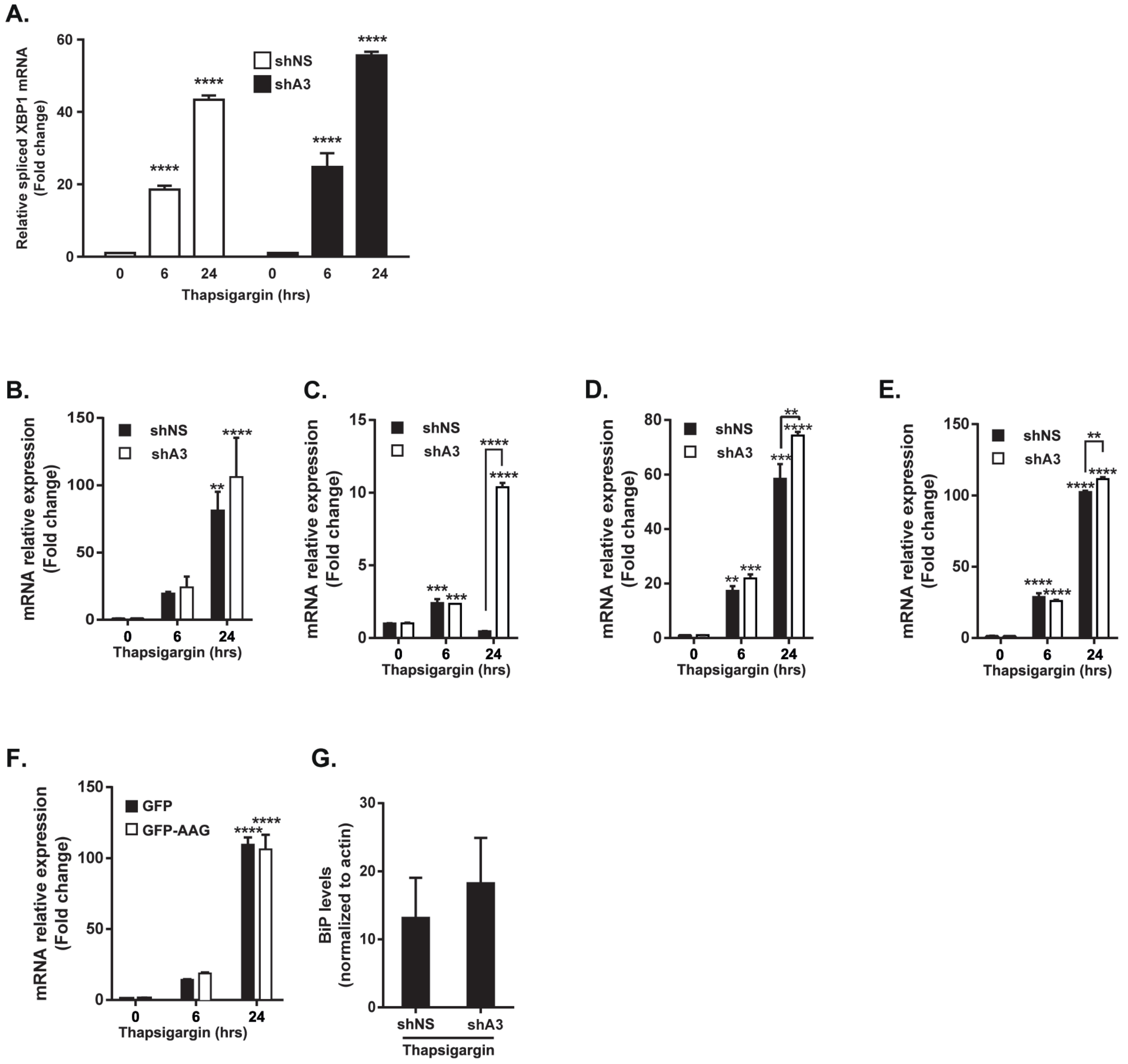


**Fig. S6. UPR transcriptional targets.** (A) XBP1s induction by thapsigargin (TG) 300nM in T98G cells treated for 6 and 24h. mRNA levels post TG of (B) BiP (C) ATF4, (D) HERPUD and (E) CHOP were analysed by qPCR. (F) BiP mRNA levels in A172 cells post TG. (G) BiP protein levels in T98G cells after TG treatment. Results indicate the mean ± S.E. of three independent experiments. ***P* <0.01, *****P* <0.0001.

#### Supplemental Figure 7

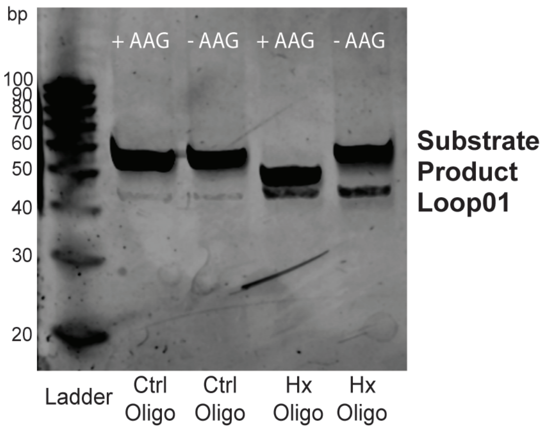


**Fig. S7** – Hypoxanthine (Hx) containing oligonucleotide (Substrate) is cleaved after incubation with recombinant hAAG and NaOH into a smaller product (Product). Loop01 signals the presence of unligated fluorescent damage-free oligo.

### Supplemental Figure 8

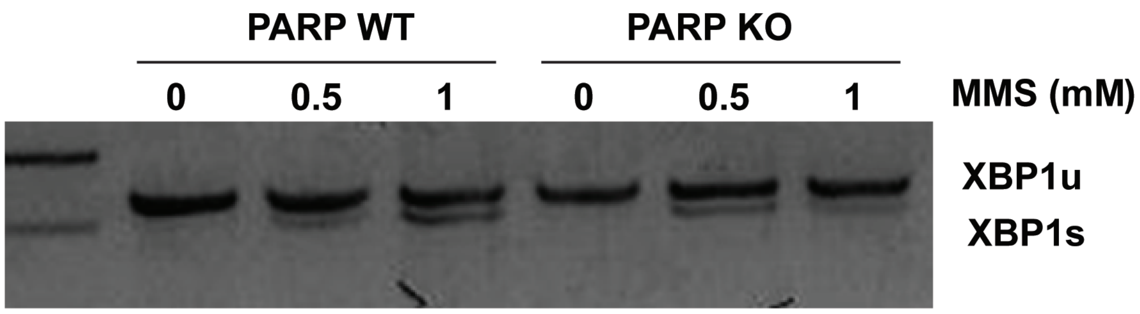


**Fig. S8** – *PARP* knock-out cells are proficient in alkylation-induced XBP1 splicing. U2OS cells, wildtype (WT) or knockout (KO) for PARP, were treated with the indicated doses of MMS for 1h in serum-free media and XBP1 splicing assessed 1h post-treatment.
